# Genome-wide CRISPR-dCas9 screens in *E. coli* identify essential genes and phage host factors

**DOI:** 10.1101/308916

**Authors:** François Rousset, Lun Cui, Elise Siouve, Florence Depardieu, David Bikard

## Abstract

High-throughput genetic screens are powerful methods to identify genes linked to a given phenotype. The catalytic null mutant of the Cas9 RNA-guided nuclease (dCas9) can be conveniently used to silence genes of interest in a method also known as CRISPRi. Here, we report a genome-wide CRISPR-dCas9 screen using a pool of ~ 92,000 sgRNAs which target random positions in the chromosome of *E. coli*. We first investigate the utility of this method for the prediction of essential genes and various unusual features in the genome of *E. coli*. We then apply the screen to discover *E. coli* genes required by phages λ, T4 and 186 to kill their host. In particular, we show that colanic acid capsule is a barrier to all three phages. Finally, cloning the library on a plasmid that can be packaged by λ enables to identify genes required for the formation of functional λ capsids. This study demonstrates the usefulness and convenience of pooled genome-wide CRISPR-dCas9 screens in bacteria in order to identify genes linked to a given phenotype.

## Introduction

The technological applications of RNA-guided nucleases derived from the Clustered Regularly Interspaced Short Palindromic Repeat (CRISPR) prokaryotic immune system (Brouns et al. 2008; Barrangou et al. 2007; Garneau et al. 2010) represents a true paradigm shift in our ability to manipulate cells at the genetic level (Barrangou and Doudna 2016). In particular, the Cas9 nuclease from type II systems can be guided by a chimeric single-guide RNA (sgRNA) which directs it to specifically cleave target sequences (Jinek et al. 2012). The ease with which these tools can be reprogrammed enables the development of powerful high-throughput screens. Recent studies have shown how an engineered CRISPR system packaged in lentiviral vectors can be used to deliver libraries of guide RNAs to human cells and create libraries of gene knockouts (Shalem et al. 2014; Wang et al. 2014). Such libraries can target most genes in the genome and were used, among other things, to identify essential genes in the human genome (Wang et al. 2015; Bertomeu et al. 2018; Evers et al. 2016; Morgens et al. 2016; Hart et al. 2015) and genetic requirements for different human viruses (Ma et al. 2015; Zhang et al. 2016; Park et al. 2017).

This approach is not directly applicable to bacteria where Cas9 cleavage leads to cell death rather than gene knockout (Bikard et al. 2014; Gomaa et al. 2014; Cui and Bikard 2016; Bikard et al. 2012; Citorik et al. 2014). The catalytic dead variant of Cas9, known as dCas9, binds specific targets without introducing a double-strand break and can conveniently be used to silence genes in bacteria (Bikard et al. 2013; Qi et al. 2013). Arrayed libraries of a few hundreds of guide RNAs have already proven useful to decipher the function of essential genes in *Bacillus subtilis* and *Streptococcus pneumoniae* (Peters et al. 2016; Liu et al. 2017). While arrayed libraries enable to access many phenotypes and easily link them to genotypes, they are cumbersome to work with and maintain. Pooled screens present the major advantage of enabling the study of much larger libraries at a low cost and with easy techniques.

We recently constructed a pooled library of ~ 92,000 sgRNAs targeting random positions in the genome of *E. coli* MG1655 with the unique requirement of a proper NGG protospacer adjacent motif (PAM) (Cui et al. 2018). This library enabled to investigate the properties of CRISPR-dCas9 screens in *E. coli* in an unbiased way. It corroborated the observations made in previous studies that dCas9 can silence gene expression following two different mechanisms: it can block transcription initiation when binding the promoter region or block transcription elongation when binding within a gene (Qi et al. 2013; Bikard et al. 2013). The strand orientation of dCas9 binding does not seem to matter when blocking transcription initiation; however binding of the guide RNA to the coding strand is required to block the running RNA polymerase. An important feature to consider when performing CRISPR-dCas9 assays is that targeting a gene in an operon will also block the expression of all downstream genes. This screen also uncovered unexpected sgRNA design rules. In particular dCas9 appears to be toxic in *E. coli* when guided by sgRNAs sharing some specific 5 nt PAM-proximal sequences. This “bad seed” effect is particularly pronounced at high dCas9 concentrations and can be alleviated by tuning dCas9 levels while maintaining strong on-target repression. Finally, this screen uncovered how 9 nt of identity between the PAM-proximal region of a guide and an off-target position in an essential or near-essential gene can be sufficient to block its expression and cause a strong fitness defect. In this study, we now analyze the results of the screen taking all these design rules into account to investigate what it can teach us about gene essentiality in *E. coli* during growth in rich medium. We show that this method can be used to confidently predict gene essentiality in most cases despite the polar effect produced by dCas9. Among other findings, we further reveal the importance of some repeated elements, identify essential genes that cannot tolerate small reductions in their expression levels, and challenge the essentiality of a few genes.

As a proof of concept of the usefulness of this approach, we then applied the screen to identify *E. coli* genes required for bacteriophage infection. The study of phage-host interactions has led to the development of powerful tools in genetics and molecular biology (Henry and Debarbieux 2012). Their study can also prove useful to understand how bacteria can mutate to become resistant, and provide insight for the design of improved phage therapies. In addition to approaches based on reverse genetics, genome-wide screens have been performed to identify host dependencies of *E. coli* phages T7 and λ using the Keio collection, an in-frame single-gene knockout strain collection (Maynard et al. 2010; Qimron et al. 2006; Baba et al. 2006). These approaches show several limitations, including the time-consuming development of such strain collections, the laborious screening process and the focus on nonessential genes only. Applied to phages λ, T4 and 186, our screen revealed various host factors including phage receptors and lipopolysaccharide (LPS) requirements. It also highlighted capsule synthesis as a shared resistance mechanism to the three phages. We finally took advantage of the ability of phage λ to package plasmids carrying a *cos* site to perform a pooled transduction assay of the library, enabling to distinguish host genes used by phage λ to lyse its host from host genes required for the production of functional λ particles.

## Results

### Screen design

In our previous work, we constructed strain LC-E75 carrying dCas9 on its chromosome under the control of an aTc-inducible promoter (Cui et al. 2018). The expression of dCas9 in this strain was fine-tuned to limit the bad seed effect while maintaining strong on-target repression. The sgRNA library was introduced into strain LC-E75 and cells were grown over 17 generations in rich medium with dCas9 induction (Fig. 1A). During this experiment, guides that reduce the cell fitness, for instance by blocking the expression of essential genes, are depleted from the library. The sgRNA library was extracted and sequenced before and after dCas9 induction. We used the number of reads as a measure of the abundance of each guide in the library and computed the log2-transformed fold change (log2FC) as a measure of fitness.

**Figure 1.**
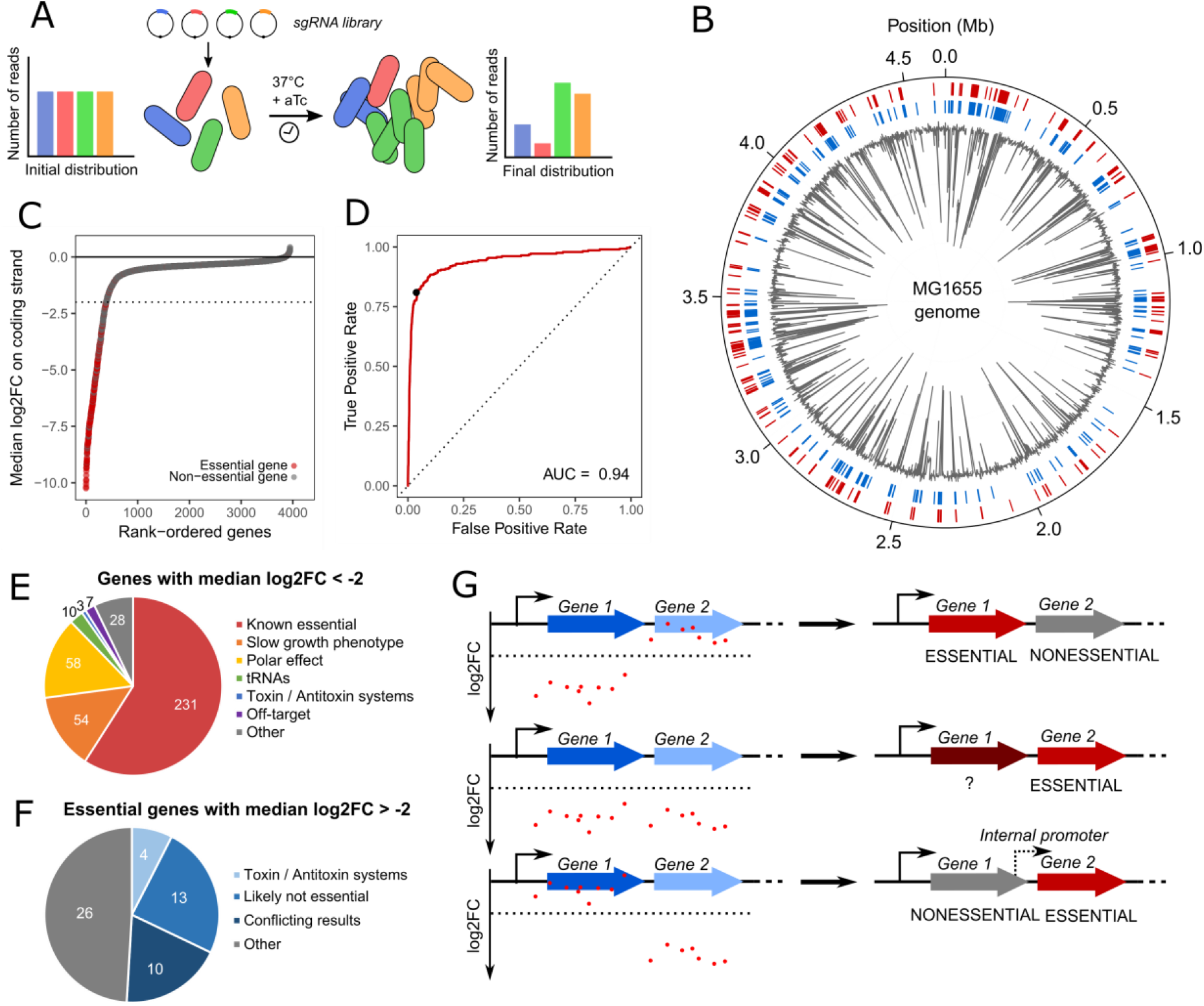
Predicting *E. coli* essential genes from the CRISPR-dCas9 screen data. (*A*) A sgRNA library was transformed into cells expressing dCas9 under the control of an aTc-inducible promoter. The distribution of sgRNAs was retrieved by deep sequencing before dCas9 induction and after 17 generations of induction. (*B*) Genome-wide visualization of the screen results. The median log2FC of sgRNAs targeting the coding strand of genes is displayed in grey. The outer red track represents genes annotated as essential in the EcoGene database while blue track shows genes selected as candidate essential in our screen (median log2FC < −2) (*C*) Genes were ranked according to the median log2FC of the sgRNAs targeting the coding strand. Genes annotated as essential in the EcoGene database are shown in red. Dashed line represents the chosen essentiality threshold (median log2FC = −2). (*D*) The median log2FC of sgRNAs targeting the coding strand was used to predict gene essentiality. Receiver Operating Characteristic (ROC) curve of the prediction model is plotted (AUC = 0.94). The black dot represents the chosen threshold. (*E*) 391 candidate essential genes were chosen with the threshold. Polar effect indicates genes located upstream essential or near-essential genes. Slow-growth phenotype indicates non-essential genes known to have a growth defect when mutated or deleted. (*F*) 53 genes annotated as essential were not classified as essential in our screen. Genes found essential only in the Keio collection but not by other sources are termed as “likely not essential”. Conflicting results indicates genes with discrepancies between various datasets. (*G*) Within an operon, different scenarios are displayed as well as their corresponding conclusion.

### Effect of guides targeting multiple positions

While most guides in our screen target unique positions in the genome, our library also includes guides targeting duplicated regions. These guides can be conveniently used to investigate the role of regions or genes present in several copies in the genome, which cannot be easily achieved by other methods. Interestingly, out of 1990 sgRNAs with multiple targets around the genome, 369 are strongly depleted (log2FC < −2). Most of these depleted guides target genes involved in translation: ribosomal RNAs, tRNAs, and the elongation factor Tu encoded by highly similar *tufA* and *tufB* genes (Supplemental Fig. S1). More surprisingly, 39 of these guides target Repetitive Extragenic Palindromic (REP) elements. Some REP elements were reported to play a regulatory role at the translational level by modulating mRNA stability (Liang et al. 2015), but their exact role remains elusive. For many of these guides, none of the target positions are in the vicinity of essential genes, suggesting that the fitness defect caused by dCas9 binding at REP sequences is not simply due to an effect on the expression of essential genes. Guides targeting several repeat regions at the same time induced a significantly stronger fitness defect than guides targeting only one repeat region. Further work will be required to understand the mechanism responsible for the fitness defect produced by REP-targeting guides, which will likely reveal interesting biology.

### Identification of essential genes

Starting from the initial library of ~ 92,000 sgRNAs targeting random positions along the chromosome of *E. coli*, we filtered sgRNAs with either a bad seed effect, an off-target, multiple targets in the genome or a low number of reads (see Methods), yielding a library of ~ 59,000 guides used to perform the analyses below (Supplemental Table 2). To investigate the usefulness of such CRISPR-dCas9 screens to identify essential genes, we first ranked genes according to the median log2FC of sgRNAs targeting the coding strand (Supplemental Table 5). Mapping our data to the genome of *E. coli* highlighted a good concordance with previously known essential genomic regions (Fig. 1B). As expected, genes with the most depleted sgRNAs included a large majority of genes annotated as essential in the EcoGene database (Zhou and Rudd 2012) (Fig. 1C). Indeed, the median log2FC value was a very good predictor of gene essentiality (Fig. 1D). Overall, a gene network analysis of the 100 top-scoring genes highlighted the main essential functions, namely ribosome assembly, peptidoglycan synthesis, DNA replication and transcription, fatty acid metabolism and tRNA metabolism (Supplemental Fig. S2).

Candidate essential genes were selected when the median log2FC of sgRNAs targeting their coding strand was lower than −2. This threshold selected 391 genes, including 231 genes annotated as essential (Fig. 1E). Among the 160 remaining genes, 54 are known to show a slow-growth phenotype when mutated or deleted and can thus be termed as near-essential. The effect of 58/160 genes can be explained by a polar effect leading to the repression of a downstream essential or near-essential gene in the same operon, such as *yejL* and *ypaB* (Fig. 2B). All in all, polar effects account for a false-positive discovery rate of essential genes of 14.8%. Another 10/160 genes are tRNA genes whose targeting leads to the silencing of several tRNAs in the same operon or are known to induce a fitness defect. In addition, 3/160 genes, *rnlB, hipB* and *ratA* are part of toxin/antitoxin systems. *rnlB* is expected to be essential (Koga et al. 2011), although it is not annotated as such in available databases for unknown reasons. Blocking the expression of *hipB* will also silence the downstream *hipA* toxin. This is expected to be toxic as degradation of HipB by the Lon protease will free HipA (Hansen et al. 2012). The antitoxin of *ratA* was not described and its regulation is poorly understood but we can make the hypothesis that guides targeting this gene also block the expression of the antitoxin. Among the 35 remaining genes, 28 remain ambiguous either because they are targeted by a very low number of sgRNAs in our library or because these genes may be essential or near-essential in our experimental conditions. Note that 5 of these genes (*aceF, lpd, dcd, hemE* and *ihfA*) were found to be essential in a recent Transposon-Directed Insertion-site Sequencing (TraDIS) screen (Goodall et al. 2018). Finally, 7 genes can be explained by off-target effects. When filtering the data to eliminate guides with off-targets, we made a compromise between the number of guides to keep in the analysis and the stringency of the filter. As a result, a few guides with off-target effects remain in the library. Since the median log2FC was used to assess gene essentiality, a gene targeted by only one or two guides might be classified as essential if one of the guides has an off-target. Two such examples are given in Supplemental Fig. S3.

**Figure 2.**
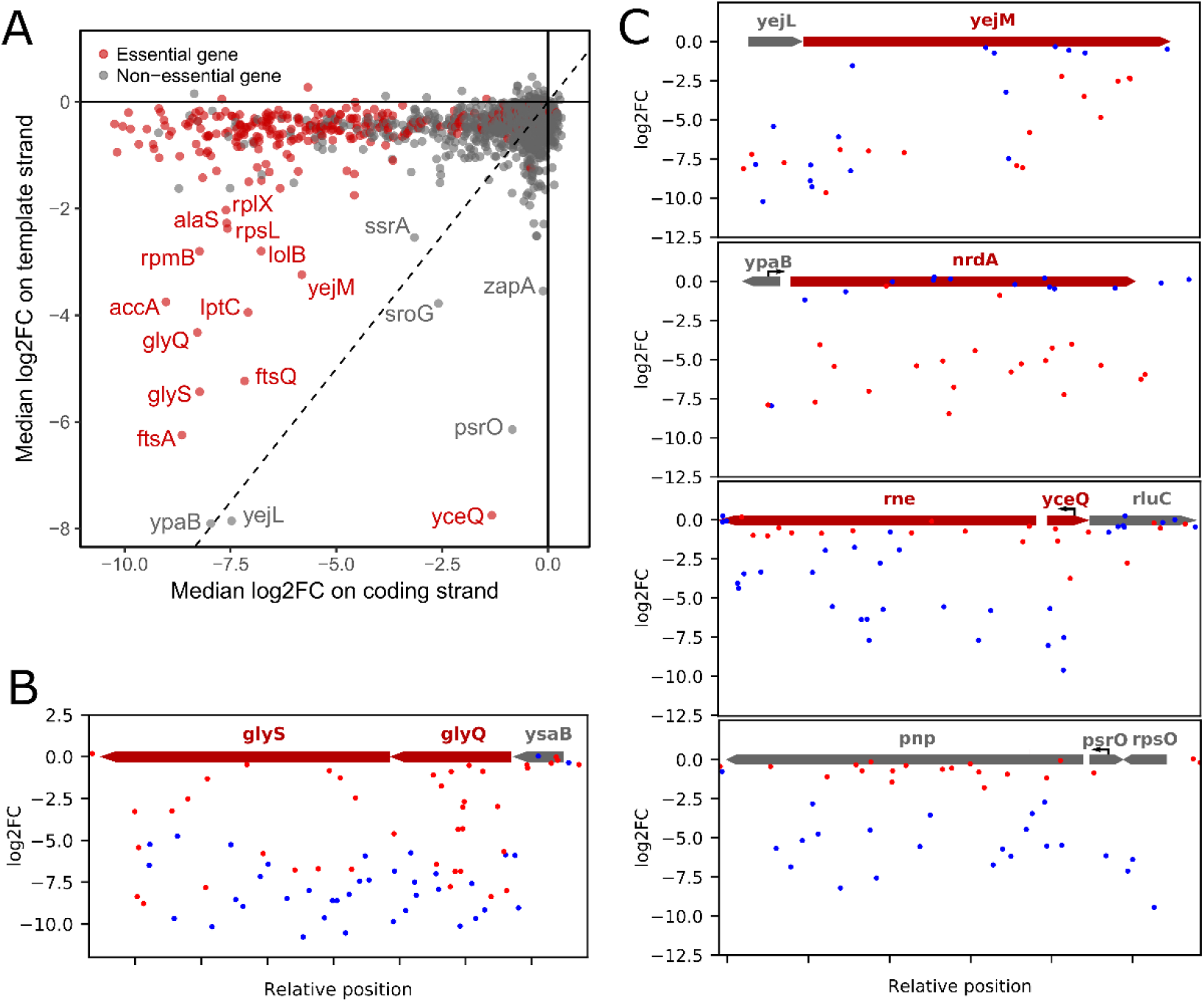
Unexpected effects highlight unusual operon architecture. (*A*) Comparison between median log2FoldChange of sgRNAs targeting the coding strand or the template strand highlights unexpected effects. (*B*) Representation of sgRNAs targeting *glyQ* and *glyS* where targeting the template strand is sufficient to induce a fitness defect. (*C*) Representation of sgRNAs targeting *yejL, ypaB, yceQ* or *psrO*. (*B,C*) sgRNAs targeting the positive or negative strand are dotted in red or blue respectively. Black arrows indicate known promoters.

In many cases, several genes with a fitness defect are adjacent within an operon. In this scenario, only the last gene with a fitness defect can confidently be classified as essential or near-essential due to polar effects. Upstream genes might or might not cause a fitness defect when silenced alone, and can be classified as potentially essential (Fig. 1G). When doing the analysis in this manner rather than gene by gene, a fitness defect can be confidently attributed to only 154 genes while 230 genes remain uncertain.

Another interesting scenario when looking at operons is when silencing a gene upstream of an essential or near-essential gene induces no fitness defect despite the theoretical polar effect. A likely explanation is that an unidentified promoter internal to the operon drives the expression of the essential gene independently of the upstream gene (Fig. 1G). We identified 7 such cases (Supplemental Fig. S4), and in all instances recent data supports the existence of an internal promoter (Thomason et al. 2015).

Conversely, some genes annotated as essential do not show a fitness defect in our screen (median log2FC > −2) (Fig. 1F, Supplemental Table 5). Out of 53 essential genes that we fail to identify, 4 genes (*chpS, mazE, yafN* and *yefM*) are part of toxin-antitoxin systems which are encoded in operons with the antitoxin gene preceding the toxin gene. By definition, an antitoxin is essential to the cell as silencing it leads the cognate toxin to kill the cell. In the four cases above, dCas9 blocks the expression of both the antitoxin and the toxin gene, which we expected to be toxic if the antitoxin is more labile. Surprisingly, unlike *hipB*, no effect on fitness was measured here. Another 13/53 genes annotated as essential in the Ecogene database were also supposed to be non-essential by other studies (Gerdes et al. 2003; Yamazaki et al. 2008; Goodall et al. 2018). We successfully replaced three of these genes, *alsK, bcsB* and *entD* by a kanamycin resistance cassette, confirming that they are indeed not essential (Supplemental Fig. S5). In 10/53 other cases, gene essentiality has also been challenged by conflicting results (Baba et al. 2006; Yamamoto et al. 2009; Goodall et al. 2018; Gerdes et al. 2003; Yamazaki et al. 2008). For the remaining 26 genes, the absence of fitness defect could be explained by an inefficient repression by dCas9, a strong robustness of the cell to low levels of the protein or could indicate genes that are actually not essential in our experimental conditions. In addition, weak repression could arise from negative regulatory feedback loops. Binding of dCas9 to genes under such control would induce an increased transcription initiation rate, ultimately leading to a weak silencing (Vigouroux et al. 2018).

### Unexpected effects on the template strand of target genes

We further investigated the effect of sgRNAs targeting the template strand of genes. Guides in this orientation should only have a moderate effect on gene expression. As expected, the vast majority of them do not produce any fitness defect (Cui et al. 2018). We compared the median log2FC of sgRNAs targeting the coding strand and the template strand (Fig. 2A). Interestingly, we observed that a set of essential genes produced a strong fitness defect regardless of the targeted strand (including *accA, alaS, ftsA, ftsQ, glyQ, glyS, lolB, lptC, rplX, rpmB, rpsL* and *yejM)*. The cell thus seems very sensitive to the expression level of these genes. Two nonessential genes, *yejL* and *ypaB*, also show a strong fitness defect with guides in both orientations, but which can in this case be explained by polar effects (Fig. 2B): *yejL* is located upstream of the essential gene *yejM*, and *ypaB* contains the promoter of the essential gene *nrdA*, whose expression is likely blocked by guides targeting *ypaB* in either orientations.

Unexpectedly, a few genes show a fitness defect when targeted on the template strand but not on the coding strand (Fig. 2A). These effects can also be explained by polar effects on nearby essential genes and shed light on atypical gene organizations. For instance, the *sroG* gene is actually a riboswitch controlling the expression of the essential gene *ribB* involved in riboflavin synthesis. One sgRNA binding to the template strand in the very beginning of the gene was strongly depleted, suggesting that it blocks transcription initiation of the *ribB* mRNA. *yceQ* is a small open reading frame of just 321bp in the promoter region of the essential RNAse E gene (*rne*) and in the opposite orientation. Guide RNAs directing dCas9 to bind the template strand of *yceQ* are thus in the good orientation to effectively block the expression of *rne* (Fig. 2B). The *rne* mRNA has indeed been described as having a long 5’ untranslated region antisens to *yceQ* which carries hairpins involved in post-transcriptional regulation (Jain and Belasco 1995). This suggests that *yceQ* is actually not essential, which is also supported by recent TraDIS data (Goodall et al. 2018). A similar case is that of *psrO* which encodes a small RNA and located in the promoter region of *pnp*. Binding of dCas9 to the template strand of *psrO* likely blocks transcription of *pnp* (Fig. 2B).

Overall, these results show the performance of CRISPRi screens to assess gene essentiality in *E. coli* but also to highlight diverse genomic organizations.

### Identification of genes providing phage resistance when silenced

To demonstrate the broad usefulness of this strategy to the study of different phenotypes, we applied the method to unveil bacterial genes required for successful phage infection, also known as host factors. Considering previous results, we now focused on sgRNAs targeting genes on the coding strand of genes only, yielding a library of ~ 22,000 sgRNAs. As a proof of concept, we used temperate phage λ whose host requirements are well documented. We also used temperate phage 186cIts (a thermosensitive strain of phage 186) and virulent phage T4 in order to compare their host requirements. As strain LC-E75 carries the *dcas9* expression cassette integrated in the *attB* site of phage 186, we constructed a new strain, FR-E01, with the same cassette integrated in the *attB* site of phage HK022 in order to avoid any interference with phage 186. Both strains expressed dCas9 at the same level (Supplemental Fig. S6).

A culture of strain FR-E01 carrying the library was grown with aTc to stationary phase allowing for dCas9 to be expressed and for the target gene products to be diluted and/or degraded. Cells were then diluted 100-fold and grown to exponential phase, still with aTc, followed by infection with phage λ, T4 or 186cIts at a MOI of 1 (Fig. 3A). During the experiment, phages lyse bacteria unless the sgRNA they carry makes the cell resistant to infection. The pool of sgRNAs was sequenced before and after 2 h of infection and a log2FC value was computed for each sgRNA as previously (Supplemental Table 3 and Fig. 3B). For each gene, a resistance score was calculated as the median log2FC of the sgRNAs targeting the coding strand (Supplemental Table 6). As expected, the highest-scoring genes correspond to phage receptors and their positive regulators (Fig. 3B-C). The phage λ receptor *lamB* (maltose outer membrane porin) (Randall-Hazelbauer and Schwartz 1973) and its regulators *malT* and *cyaA* respectively ranked 2^nd^, 1^st^ and 6^th^ after infection by λ phage. The 3^rd^ place was occupied by gene *malK* which is expressed upstream of *lamB* in an operon structure. Guides that block the expression of this gene therefore also block the expression of *lamB*, explaining its high ranking although it is not known to participate in the infection process. Phage T4 can either use the outer membrane porin OmpC as a receptor or the core-lipid A region of the LPS that includes the heptose region (Yu and Mizushima 1982). Gene *ompC* and its regulator *ompR* respectively ranked 1^st^ and 2^nd^ after T4 infection, while genes *rfaD* and *waaF* respectively involved in the synthesis of glycerol-D-manno-heptose and its addition to the LPS, ranked 5^th^ and 8^th^. Considering that the receptor for phage 186 is not clearly established, we used this approach to identify candidate host components. Genes with the highest resistance scores to 186 did not include any surface protein but did include *waa* genes involved in the LPS synthesis pathway (Fig. 3C). Some of them, including *rfaD-waaF*, were also found to affect phage λ infection in our screen and are known to be associated with lower levels of LamB (Randall 1975), but the LPS requirement was stronger for phage 186. To validate this observation, we performed infection time courses with a strain deleted for *waaJ*, one of the last genes in the LPS biosynthesis pathway, in the presence of each of the three phages. The absence of gene *waaJ* induced a strong resistance to 186 but not to λ and T4 (Fig. 3D) suggesting that the outer core of the K-12 strain LPS is required for adhesion of phage 186. Taken together, these observations validate the ability of our method to identify phage receptors which are the most common bacterial components giving rise to phage resistance.

**Figure 3.**
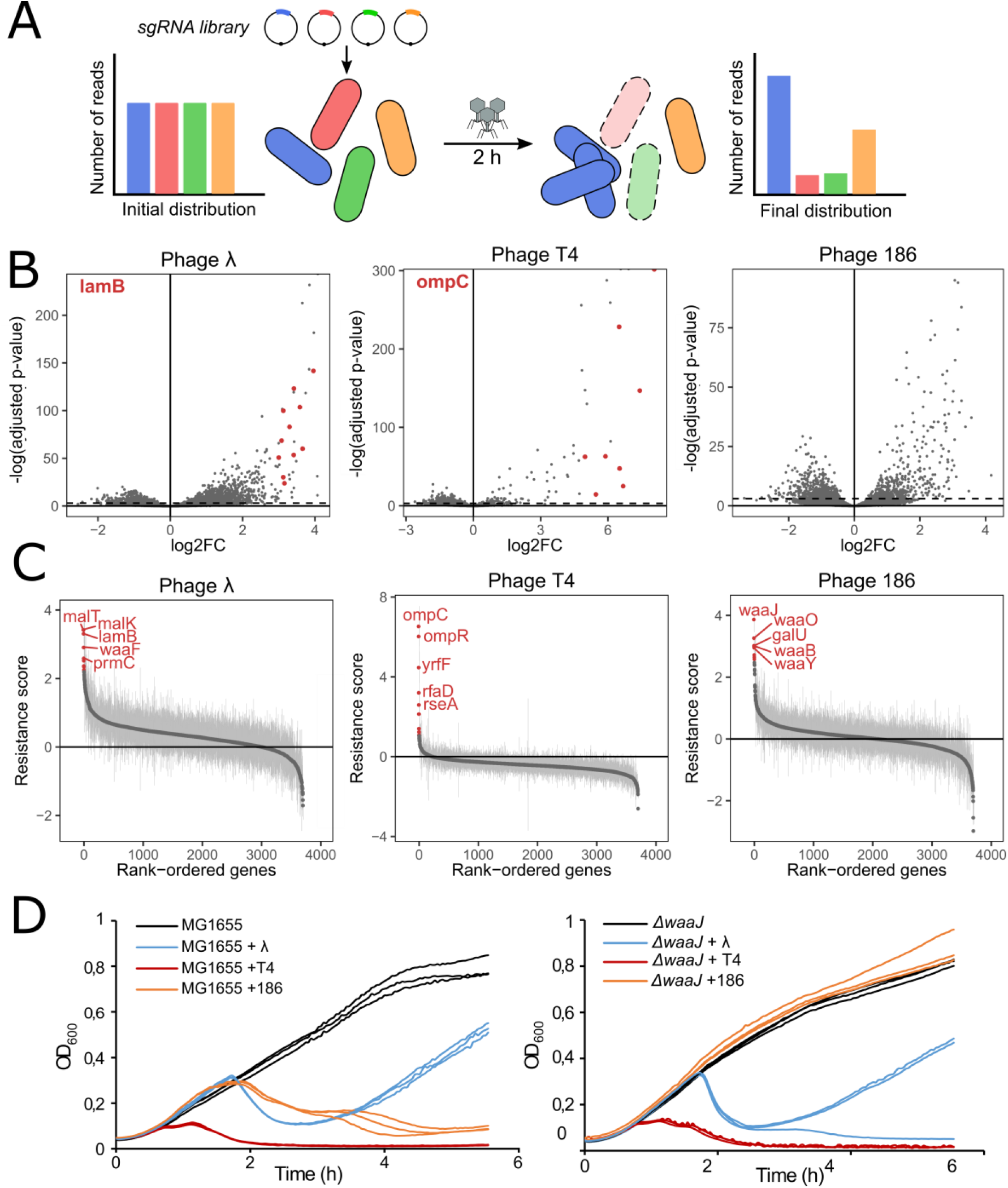
The CRISPRi screen reveals phage receptors and LPS requirements of phages λ, T4 and 186. (*A*) Overview of the experimental procedure. dCas9 expression was induced for ~ 8h before infection by λ, T4 or 186 at a MOI of 1. The distribution of sgRNA was retrieved before and after infection by sequencing. (*B*) The log2FoldChange was computed for each sgRNA. For phages λ and T4, sgRNAs targeting the phage receptor (respectively LamB and OmpC) are highlighted in red. Dashed line represents adjusted p-value = 10^−3^. (*C*) For each gene, a resistance score was calculated as the median log2FoldChange of sgRNAs targeting the coding strand. 5 top-scoring genes are highlighted in red in each panel. (*D*) Infection time courses of MG1655 (top panel) or Keio collection strain *ΔwaaJ* without phage or after infection by λ, T4 or 186 (MOI ~ 0.01).

Further comparison between phages λ, T4 and 186 revealed closer host requirements between the two temperate phages, λ and 186, than between λ and T4 or between 186 and T4 (Fig. 4A). We selected genes with a resistance score of at least 20% of the maximum score for each phage (Fig. 4B). This threshold selected only 5 genes for phage T4 while it selected 367 genes for phage λ and 88 for phage 186, suggesting that T4 relies on very few host components. Genes selected after λ infection included the *rnpA* and *rnt* RNAses as well as genes involved in two tRNA modification pathways: uridine 2-thiolation (*tusA, tusBCD, tusE* and *mnmA*) and the essential N^6^-threonylcarbamoyladenosine modification (*tsaB, tsaC, tsaD* and *tsaE)*. Note that uridine 2-thiolation has been previously described to be necessary for the proper translation of λ proteins gpG and gpGT involved in tail assembly through its impact on programmed ribosomal slippage (Maynard et al. 2010, 2012). We can hypothesize that the essential N^6^-threonylcarbamoyladenosine modification of tRNAs is also necessary for the proper translation of λ proteins. Interestingly, genes required for both λ and 186 infection included *prmC* involved in translation termination, genes *dnaKJ* known to be required for DNA replication of λ (Yochem et al. 1978), as well as lysogenization regulator *hflD*. During λ infection, HflD facilitates degradation of repressor CII, thus avoiding lysogeny (Kihara et al. 2001). This result suggests that HflD could also regulate lysogeny in phage 186.

**Figure 4.**
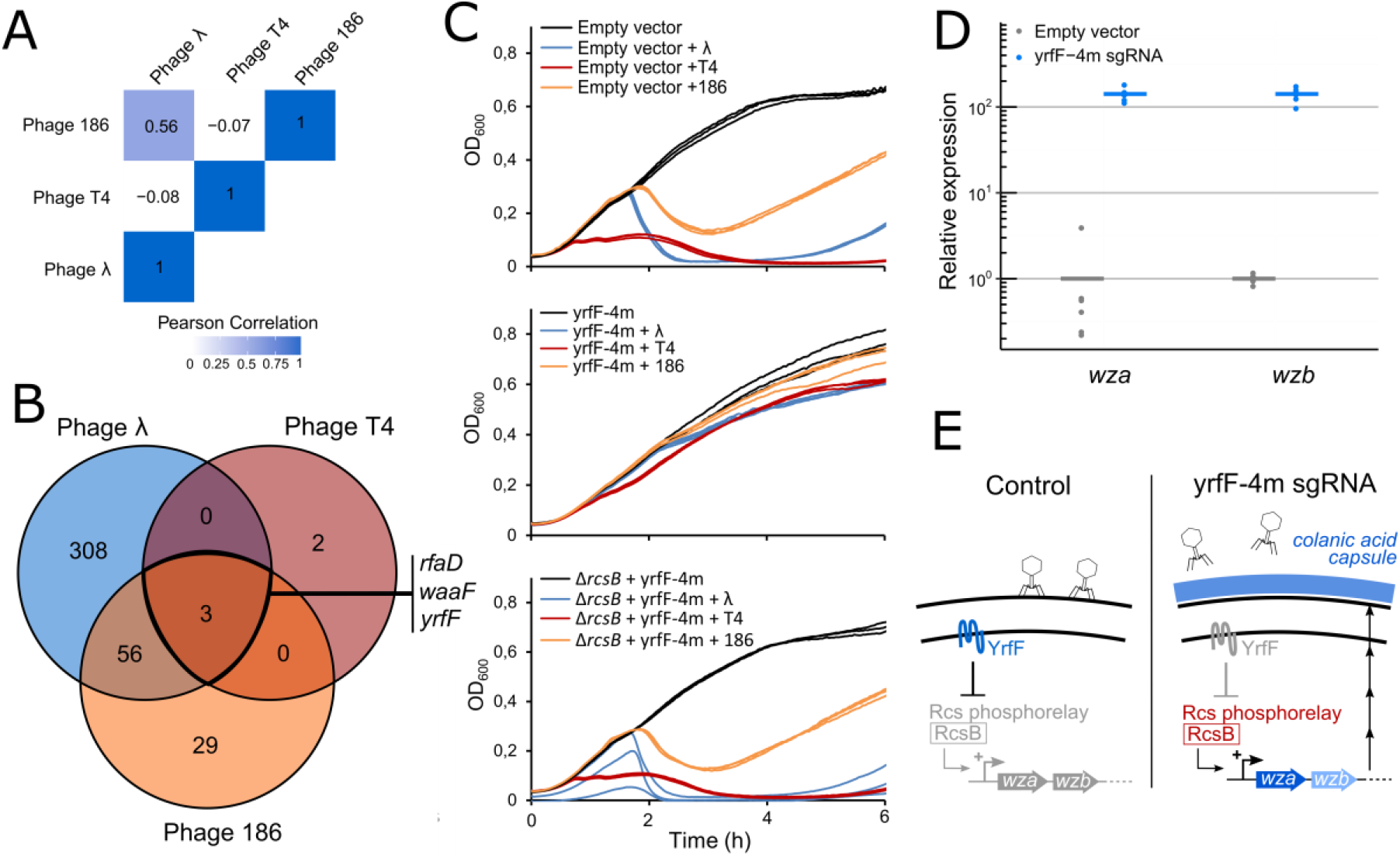
Colanic acid capsule synthesis is a common resistance mechanism to phages λ, T4 and 186. (*A*) Pearson correlation coefficient of resistance scores between phages λ, T4 and 186-cl. (*B*) For each phage, genes with a resistance score greater than 20% of the maximum score were selected and compared in a Venn diagram. (*C*) Infection time courses of FR-E01 carrying an empty vector, FR-E01 expressing *yrfF* sgRNA with 4 mismatches (yrfF-4m) or FR-E0MrcsB expressing yrfF-4m without infection or after infection by λ, T4 or 186-cl (MOI ~ 0.01). Experiments were performed in triplicates. (*D*) Expression of genes *wza* and *wzb* involved in colanic acid capsule synthesis in strain FR-E01 carrying an empty vector or expressing yrfF-4m sgRNA, as measured by RT-qPCR. Results are shown for 3 biological replicates and 2 technical replicates. (*E*) Schematic of colanic acid capsule-mediated phage resistance. YrfF inhibits the Rcs phosphorelay. Inhibition of YrfF expression activates the Rcs phosphorelay which triggers the expression of genes involved in colanic acid capsule synthesis.

Three genes with a high resistance score to all 3 phages were identified (Fig. 4B): *rfaD* and *waaF* discussed above for their involvement in LPS synthesis, and *yrfF*, an essential gene indirectly linked to capsule synthesis. YrfF inhibits the Rcs signaling pathway by an unknown mechanism (Clarke 2010). This signaling pathway senses and responds to damage to the membrane or to the peptidoglycan by activating genes involved in colanic acid capsule synthesis (Gottesman and Stout 1991; Stout 1994), suggesting that *yrfF* silencing induces capsule synthesis. Since this gene is essential, it cannot be deleted or repressed by a fully matched sgRNA without killing the cell. We thus reduced its expression to intermediate levels using a sgRNA bearing 4 mismatches at the 5’-end (yrfF-4m) (Bikard et al. 2013; Vigouroux et al. 2018). The resulting strain only showed a slight growth defect and became resistant to all three phages (Fig. 4C). Deletion of gene *rcsB*, a central actor of the Rcs pathway, restored sensitivity to the 3 phages in this strain, showing that the resistance phenotype that results from *yrfF* silencing is mediated by the Rcs pathway. We further confirmed this by showing that silencing *yrfF* induces a ~ 140-fold increase in the expression of the *wza-wzb* operon directly involved in colanic acid capsule synthesis (Fig. 4D-E). Interestingly, deletion of *rfaD* or *waaF* (but not other genes of the LPS biosynthesis pathway) was shown to induce a mucoid phenotype through the synthesis of a capsule (Joloba et al. 2004). This suggest that they might provide resistance to all three phages through this pathway rather than through their role in LPS synthesis. Consistently with this hypothesis, silencing genes upstream of *rfaD* and *waaF* in the LPS biosynthesis pathway did not provide resistance to T4 in our screen.

### Identification of genes involved in later steps of λ infection

The screen performed here is thus a powerful method to identify genes required by phages to kill the cell. However, this first strategy cannot identify genes necessary for the synthesis of functional phage capsids if blocking the expression of these genes does not prevent the phage from killing the bacteria. One can expect that this will be the case of any host gene involved in late stages of the infectious cycle when the host cell is already doomed. In order to get better insights into the genes affecting the production of functional phages, we implemented a second step focusing on phage λ (Fig. 5A). The vector carrying the sgRNA library was designed to carry a λ packaging site (*cos* site), thus forming a cosmid (Cronan 2013). Upon infection, phage λ is thus able to package the plasmid carrying the guide RNA only if a functional capsid was produced (Supplemental Fig. S7). After infection by phage λ, the cell lysate containing a mixture of λ and cosmid particles was used to transduce strain MG1655::λ and thus recover guides in the library that do not affect the infection process. The distribution of sgRNAs recovered after transduction was compared to the initial pool to identify depleted sgRNAs corresponding to bacterial genes required for the production of functional capsids (Supplemental Table 4). Interestingly, 45 depleted guides in our library have a perfect match in the λ phage genome, mostly in the late operon (Supplemental Fig. S8). These guides were not found to block cell lysis in our first screen as they likely act too late in the phage cycle, but are able to limit the number of particle produced.

**Figure 5.**
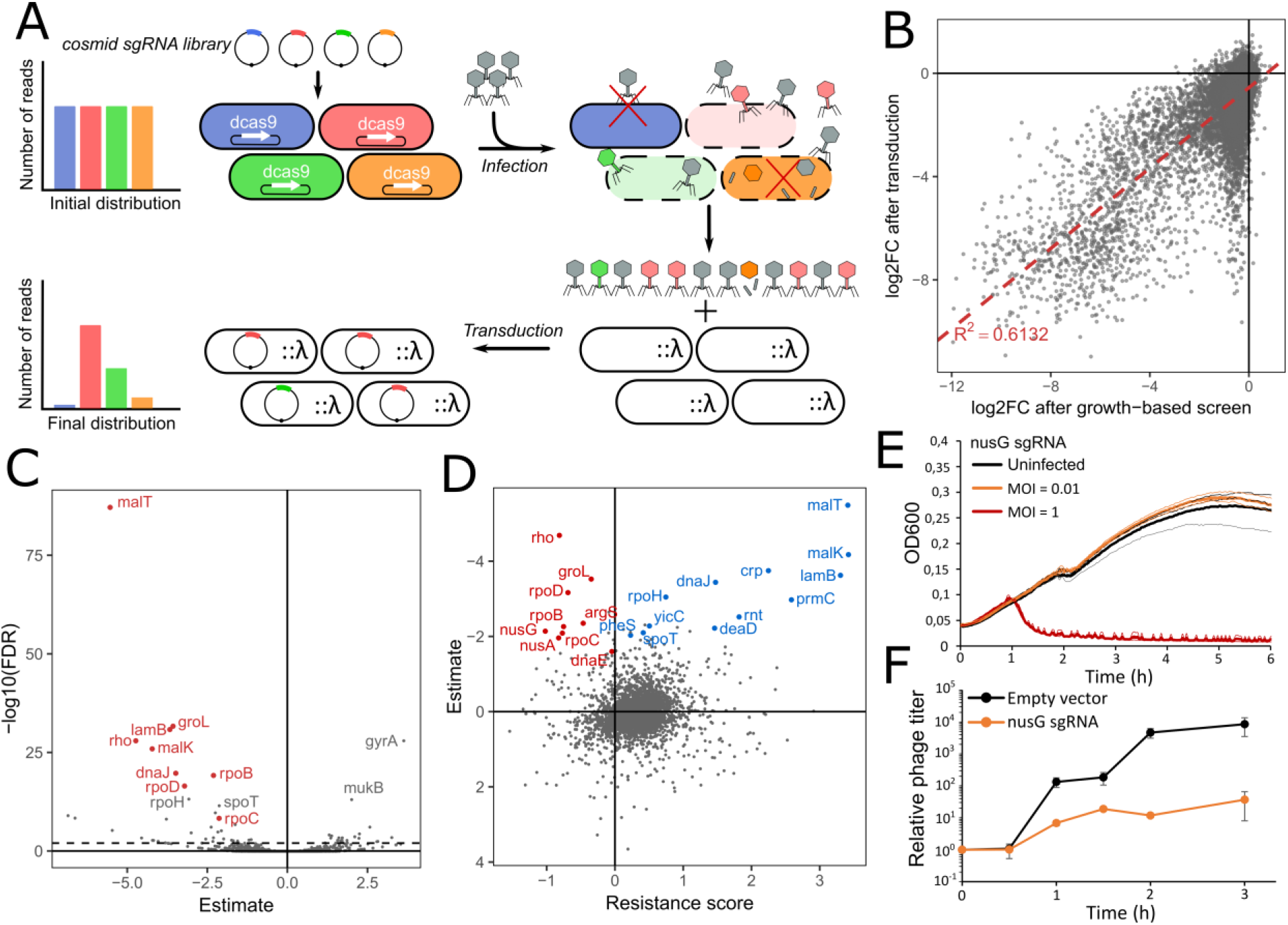
Transduction of the library reveals host genes necessary for the production of functional capsids. (*A*) Overview of the experimental system to screen for λ host factors. Strain FR-E01 expressing dCas9 under aTc-inducible promoter and carrying the sgRNA library on a cosmid was infected by λ. The library can be packaged into capsids when the sgRNA does not disrupt phage propagation. After transduction into a lysogenic strain, the distribution of sgRNAs is measured by sequencing. (*B*) log2FC values after transduction are strongly correlated to the effect of guides on cell growth (ANOVA, F = 27263, df= 17198, p < 10^−16^). (*C*) For each gene, a generalized linear model was built to explain log2FoldChange after transduction while taking the effect on cell growth into account. Estimate and corrected p-value (FDR) are shown for each gene. Genes previously described as host factors are shown in red. (*D*) Comparison of results after infection and after transduction. Genes that significantly decrease the production of functional phages (FDR < 0.001) are highlighted in red or blue when they show a negative or positive resistance score respectively. (*E*) Growth of strain FR-E01 carrying a sgRNA targeting *nusG* without infection or after infection by λ at MOI = 0.01 or MOI = 1. (*F*) Phage titer during infection of strain FR-E01 carrying an empty vector or expressing a sgRNA targeting *nusG* (MOI = 0.01). Error bars show standard deviation of three independent experiments.

When comparing our data with the essentiality data from our previous screen, we observed a strong correlation between the effect of guides on the cell fitness and log2FC after transduction (Fig. 5B). One might indeed expect that genes whose repression slows down cell growth will also slow down phage production, leading to fewer particles being released after 2h of infection. In order to identify genes which disproportionately affect phage production over cell growth, we performed statistical analyses taking the effect on growth into account (see Methods, Supplemental Table 7). We identified 57 genes which significantly decrease the amount of transduced particles when repressed (FDR < 0.05) (Fig. 5C). These genes could be separated into two groups according to their resistance score (Fig. 5D): genes with a positive resistance score provided resistance to lysis when silenced, while genes with a negative resistance score led to an increased sensitivity to the phage but a reduced production of functional phage particles. The first group includes genes involved in the expression of LamB (*lamB, malK, malT, crp, cyaA*) as well as other genes identified in the first step, such as *rpoH* and *dnaJ, prmC, rnt* and *rnpA*. The second group includes genes that could not be identified in the first step of our screen based on their resistance score as they decrease or have little effect on cell survival to λ infection. These genes include well-characterized genes involved in the transcription of the λ genome: RNA polymerase subunits β *rpoB* and β’ *rpoC*, RNA polymerase σ^70^ factor *rpoD* and transcription termination factors *rho, nusA* and *nusG* (Li et al. 1992; Gottesman et al. 1980; Friedman 1992; Ghysen and Pironio 1972). These factors interact with λ protein N to avoid transcription termination at terminator sites present on its genome, thus allowing the expression of downstream genes in a process called transcription antitermination (Gottesman et al. 1980). The screen also identified *groL* encoding the GroEL chaperonin involved in the assembly of the λ head (Sternberg 1973b; Georgopoulos et al. 1973; Sternberg 1973a). Its transcription is controlled by the heat shock-specific RNA polymerase σ^32^ factor *rpoH* found in the first group (Chuang and Blattner 1993). Genes with little effect on survival after λ infection include DNA polymerase III subunit α encoded by *dnaE*, as well as many genes encoding aminoacyl-tRNA synthetases, namely *argS, pheS, pheT, leuS, hisS, proS, trpS, valS, cysS*, and *metG*, suggesting that these essential elements are a limiting resource for the production of phage particles.

This analysis highlights the fact that silencing some genes inhibits the formation of phage particles while leaving the cell susceptible to lysis by the phage. To confirm this observation, infection time courses and phage titer measurements were performed in strain FR-E01 expressing a sgRNA targeting *nusG* which has a negative resistance score (Fig. 5E). Infection at a MOI of 1 led to a complete lysis of the cell population showing that silencing *nusG* does not provide resistance to infection. However, infection at MOI = 0.01 did not lead to lysis of the population, presumably because the first infected cells did not release enough active phages to lyse the rest of the population. Measuring phage concentration during infection indeed showed a ~ 10-fold reduction in burst size when *nusG* is silenced (Fig. 5F).

## Discussion

This work establishes a proof of concept of the performance of genome-wide CRISPRi screens in bacteria for the identification of essential genes and phage host factors. When series of genes in operons are found to have an effect, it is theoretically only possible to conclude that the last gene in the series is essential. A gene by gene analysis of our data was nonetheless able to reliably predict gene essentiality with a false positive discovery rate due to polar effects of only 14.8%. This surprisingly good performance is due to the fact that essential genes tend to cluster in the same operons. Yet, this is not always the case and polar effects represent a limitation of what can be done with CRISPRi screens: one can easily identify operons of interest but a detailed understanding of the involvement of each gene requires further investigation with other techniques. Another limitation of CRISPRi is the heterogeneity in sgRNA repression efficiency. Efforts have been made in Eukaryotic systems to predict sgRNA activity (Kim et al. 2018; Chuai et al. 2017; Doench et al. 2016; Chari et al. 2017; Labuhn et al. 2018) but such models remain to be developed in bacteria in order to improve the quality of pooled CRISPR-dCas9 screens.

It is interesting to discuss the advantages and drawbacks of CRISPR-dCas9 screens in comparison with other genome-wide approaches. Arrayed collections of gene deletions have also proven extremely useful for the study of the few bacterial species where they have been constructed. The most commonly used is the *E. coli* Keio collection (Baba et al. 2006; Yamamoto et al. 2009) which is used as a reference for the classification of *E. coli* genes as essential, and which has been used in many different types of screens, including for the identification of phage host factors (Maynard et al. 2010; Qimron et al. 2006). However, the development of such strain collections and the screening process are laborious and focused on nonessential genes only. The Keio collection is also known to contain a few errors arising from the failure to obtain some gene knockouts for other reasons than gene essentiality, or from the fact that some genes were duplicated during the construction of the collection (Yamamoto et al. 2009). Transposon-based strategies such as Tn-seq are commonly employed in bacteria. With these approaches, false negatives may arise when the insertion of the transposon does not disrupt the functionality of the protein, or when only a small region of a gene is essential. False positives might also arise from biases in the number of transposon insertions along the genome. Consequently, recent results highlighted the need to combine statistical analysis with manual curation in transposon-based screens (Goodall et al. 2018). Finally, both methods can cause polar effects on the expression of adjacent genes which can be hard to predict as opposed to the predictable effects of dCas9 binding on the expression of downstream genes.

Overall, CRISPRi screens present several key advantages: (i) The expression of dCas9 is inducible, which enables to maintain guides targeting essential genes in the library, allowing their study; (ii) Duplicated regions can easily be studied since sgRNAs can be designed to target identical sequences present in several copies along the genome; (iii) Intermediate repression levels can be obtained either by targeting the template strand of genes or by using sgRNAs with a variable number of mismatches (Vigouroux et al. 2018; Bikard et al. 2013); (iv) libraries can be rationally designed to target specific locations or subsets of genes, as opposed to the random insertion of transposons. (v) Library sequencing is straightforward compared to the tedious process of extracting transposon junctions from genomic DNA.

In this study, we used these key advantages to investigate essential genes and other interesting features of the *E. coli* genome. Overall, 81.3 % (231/284) of the genes previously annotated as essential and covered in our study were correctly identified as essential. A large part of the genes that we failed to identify might not actually be essential. Indeed, our screen showed some discrepancies with annotated databases, particularly with the Keio collection. We experimentally confirmed that some genes were wrongly annotated as essential, corroborating other recent work (Goodall et al. 2018). Nonetheless, our screen failed to identify a few well-known essential genes. This might be due to weak dCas9 repression, strong robustness of the cell to low protein levels or negative feedback loops. Note also that the design of the library being random, some genes only have a small number of guides, reducing the likelihood to correctly identify them. This will easily be corrected in future screens by a rational library design.

In addition to the simple investigation of gene essentiality, our results highlighted the importance of REP elements and the usefulness of CRISPRi to investigate such repeated loci. Targeting the template strand of genes further enabled us to identify essential genes which cannot tolerate small reductions of their expression level. Proteins encoded by these genes could be promising antibiotic targets as a partial inhibition of their activity is likely lethal. Our screen also highlighted some atypical genomic organizations such as internal promoters within operons or nonessential genes antisense to neighboring essential genes.

Once constructed, a guide RNA library can be conveniently used to screen other phenotypes of interest. We thus applied the method to identify *E. coli* genes required for phages to kill their host. Among the host factors identified, phage receptors and other genes involved in phage adhesion had the strongest effect. Interestingly, some host factors were common between λ and 186 suggesting some similarities between their infectious cycles. Phage T4 relied on fewer host genes than λ and 186, which is consistent with the fact that T4 carries most of the components required for its replication (Miller et al. 2003). Accordingly, *E. coli* was only reported to become resistant to phage T4 by acquiring mutations that block its adsorption (Lenski 1988), whereas *E. coli* can also become resistant to λ by acquiring mutations in intracellular components (Friedman 1992). Our cosmid-based transduction assay enabled a more in-depth study of λ infection and identified genes which did not provide phage resistance when silenced. Note that our screen failed to identify a few previously described host factors for λ infection (Maynard et al. 2010), likely for the same reasons that we failed to identify a few essential genes.

A notable result of our screen is the identification of colanic acid capsule synthesis as a shared resistance mechanism to phages λ, 186 and T4. The capsule of *E. coli* has been shown to protect against external aggressions such as antibiotics or desiccation and help to evade the host immune system (Rendueles et al. 2017). While previous studies have reported that bacterial capsule can also provide resistance to several phages (Kim et al. 2015; Scholl et al. 2005; Paynter and Bungay 1970)., this is rarely seen as the main function of the capsule Our findings argue for a broader role of colanic acid capsule synthesis as a defense mechanism against a wide range of phages, acting as a physical barrier by masking phage receptors on the cell surface. Accordingly, some phages have evolved the capacity to degrade capsule through the action of enzymes such as endosialidases which are frequently carried by the tail fibers of caudovirales (Bull et al. 2010; Cornelissen et al. 2012; Scholl et al. 2001; Kim et al. 2015).

Genome-wide CRISPR screens have already demonstrated their usefulness in Eukaryotic systems. This study should now set the stage for their broader adoption as a powerful tool in bacterial genetics.

## Methods

### Bacterial strains and media

*E. coli* strains were grown in Luria-Bertani (LB) broth. LB-Agar 1.5% was used as a solid medium. Antibiotics were used in standard concentrations (50 μg / mL kanamycin, 100 μg / mL carbenicillin). *E. coli* DH5α was used as a cloning strain and *E. coli* K12 MG1655 was used for screening experiments.

### Phage strains and stocks

Phages λ and 186cI-ts were propagated from lysogenic *E. coli* strain while T4 was propagated from liquid stocks. Overnight cultures of MG1655::λ and C600::186cIts were harvested and the supernatant was filtered (0.22 μm). All the liquid stocks were further propagated in MG1655 grown in LB supplemented with maltose 0.2% and CaCl_2_ 5 mM at a multiplicity of infection (MOI) of 1. Phage titers were measured by spotting 2 μL-drops of 10-fold serial dilutions in λ dilution buffer (TrisHCl 20 mM, NaCl 0.1M and MgSO4 10 mM) on a bacterial lawn of *E. coli* MG1655 in LB-agar (0.5%) containing 5 mM CaCl_2_.

### *E. coli* strain construction

Strain LC-E75 used for the essentiality screen was described in our previous study (Cui et al. 2018). This strain expresses an optimized *dcas9* cassette under the control of a ptet promoter integrated at the phage 186 attB site. In order to study phage 186 with our screen, a new strain FR-E01 was constructed with the same cassette integrated at the HK022 attB site to avoid any interference. A high-copy pOSIP backbone (St-Pierre et al. 2013) containing the HK022 integrase, HK022 *attB* site and a Kanamycin-resistance cassette was digested by EcoRI and PstI. A fragment containing *dcas9* gene under the control of an aTc-inducible promoter (ptet) was amplified by PCR with primers LC124/LC125 to introduce homology regions with the backbone. The 2 fragments were assembled by the Gibson method (Gibson et al. 2009). The resulting vector was electroporated into strain MG1655. The backbone containing the HK022 integrase and a Kan^R^ selection marker was then removed using the pE-FLP (Amp^R^) plasmid able to recombine FRT sites flanking the backbone. pE-FLP was then cured through serial restreaks on LB-Agar, yielding strain FR-E01. Primers used for cloning are listed in Supplemental Table 1.

### CRISPRi library design and assembly

A library of ~ 92,000 sgRNAs was designed previously (Cui et al. 2018). These sgRNAs target 20-nt regions adjacent to NGG sites in *E. coli* K-12 MG1655 and were chosen randomly among the total pool of possible sgRNAs in this strain. Briefly, the library was generated through on-chip oligo synthesis (CustomArray) and assembled into the psgRNAcos backbone using the Gibson method (Gibson et al. 2009). The resulting plasmid library DNA was transferred to strains LC-E75 and FR-E01 by electroporation yielding > 10^8^ colonies, thus ensuring a ~1000X coverage of the library. Primers used for cloning are listed in Supplemental Table 1.

### High-throughput screens

Growth-based screen data was obtained from our previous study (Cui et al. 2018). This screen was performed over 17 generations in triplicates from independent aliquots of the library generated from 3 independent transformations into strain LC-E75.

The phage screen was performed in triplicates as follows: FR-E01 was grown at 37°C from 1 mL aliquots stored at −80 °C into 500 mL LB. At OD_600_ = 0.2, dCas9 expression was induced by addition of 1 nM aTc to trigger the silencing of the target genes. The culture was grown to stationary phase (OD_600_ = 2) and diluted 100-fold in LB containing 1 nM aTc, 0.2 % Maltose and 5 mM CaCl_2_. At OD_600_ = 0.4, 20 mL of the culture was sampled and the library was extracted by miniprep (Nucleospin Plasmid, Macherey-Nage) to obtain the sgRNA distribution before infection. The culture was then infected with 1 mL of λ, T4 or 186cI-ts stocks (10^7^ pfu / μL) to reach a MOI of 1. After 2 h at 37°C, the cultures were harvested (7 min – 4,000 g). Pellets were washed twice in PBS and the library was extracted by miniprep to obtain the sgRNA distribution after infection. For phage λ, the supernatant containing a mixture of λ and packaged library was collected and filtered (0.22 μm). Transduction was performed with 25 mL of lysate and 75 mL of stationary phase culture of MG1655 carrying λ prophage (providing resistance to super-infection) grown in LB supplemented with 0.2 % maltose. After 30 min at 37°C, cells were harvested and resuspended in 1 mL LB and transduced cells were selected on 4 12×12 cm Petri dishes containing kanamycin. After 4h at 37°C, nascent colonies were washed and harvested in 5 mL LB-Kan. Each sample was split into 6 minipreps to obtain the final sgRNA distribution.

### Illumina sample preparation and sequencing

Library sequencing was performed as described previously (Cui et al. 2018). Briefly, a customized Illumina sequencing method was designed to avoid problems arising from low library diversity when sequencing PCR products. Two PCR reactions were used to generate the sequencing library with primers listed in Supplemental Table 1. The 1st PCR adds the 1st index. The 2nd PCR adds the 2nd index and flow cells attachment sequences. Sequencing is then performed using primer LC609 as a custom read 1 primer. Custom index primers were also used: LC499 reads index 1 and LC610 reads index 2. Sequencing was performed on a NextSeq 500 benchtop sequencer. The first 2 cycles which read bases common to all clusters were set as dark cycles, followed by 20 cycles to read the guide RNA. Using this strategy, we obtained on average 17 million and 4.6 million reads per experimental sample for growth-based screen in LC-E75 and for phage screen in FR-E01 respectively.

### Data analysis

Samples were retrieved from pooled sequencing data according to the corresponding pairs of indexes. For essentiality analysis, guides were filtered for potential off-target effect by discarding guides whose 9 PAM-proximal bases have a perfect match next to an NGG PAM in the promoter region of a gene, or when the 11 PAM-proximal bases have a perfect match next to an NGG PAM allowing binding to the coding strand of a gene. Guides were also filtered to remove the 10 strongest bad seeds (AGGAA, TAGGA, ACCCA, TTGGA, TATAG, GAGGC, AAAGG, GGGAT, TAGAC and GTCCT) described in our previous study (Cui et al. 2018). Guides with less than 20 reads in total were discarded. Statistical analysis was performed from count data using the DESeq2 package (Love et al. 2014) in R. This package allows comparison of expression data using negative binomial generalized linear models. Read counts were normalized to a non-targeting control guide RNA present in the library. A paired analysis was performed to compare each sample to its initial condition. The log2FoldChange (log2FC) value represents the enrichment or depletion of each sgRNA. For the phage screen, guides targeting the template strand of genes or outside of genes were excluded from the analysis, yielding a library of ~ 22,000 sgRNAs. The lists of all sgRNAs with computed log2FC values after the growth-based screen, after phage screens and after transduction assay are provided as Supplemental Tables 2, 3 and 4 respectively. For each gene, the median log2FC value of the sgRNAs targeting the coding strand was used for ranking (Supplemental Tables 5 and 6). For the λ transduction experiment, a linear model was built to predict log2FC as a function of the log2FC values obtained from the growth-based screen and a Boolean parameter considering or not the gene as a host factor. A given gene was regarded as a hit when the Boolean parameter significantly improved the model after correction for multiple testing (FDR < 0.05). Genes whose silencing decreases the production of functional capsids have a negative estimate after transduction. A list of estimates and FDR for each gene is provided as Supplemental Table 7.

### Infection dynamics

sgRNAs were cloned into a psgRNAcos backbone through Golden Gate assembly using BsaI (Engler et al. 2008) and were electroporated into strain FR-E01. For essential gene *yrfF*, 4 mismatches were inserted at the 5’-end of the sgRNA to decrease expression of the target gene to intermediate levels. Constructions were validated by Sanger sequencing. The list of individual sgRNAs is provided in Supplemental Table 8. Strains were grown overnight and diluted 100-fold in LB medium containing 0.2% maltose, 1 nM aTc, 5 mM CaCl_2_ and kanamycin. At OD_600_ = 0.4, 90 μL of cultures were mixed in 96-well plates with 100 μL of fresh medium and 10 μL of a 8.10^4^ pfu/μL stock of the appropriate phage (MOI ~ 0.01). Infection dynamics were monitored in three replicates on an Infinite M200Pro (Tecan) at 37°C with shaking for 8h. Strain Δ*waaJ* was obtained from the Keio collection and infection dynamics were performed identically.

### Gene deletions

Plasmid pKOBEG-A carrying the λ-red system components under the control of an arabinose-inducible promoter was transformed into strain MG1655. A kanamycin resistance cassette was amplified from plasmid pKD4 with primers introducing homology regions with sequences flanking genes *alsK, bcsB* or *entD*. Electrocompetent MG1655-pKOBEG-A cells were prepared with arabinose before transformation of the DNA fragments (1 μg). Recovery and overnight incubation were performed at 30°C with arabinose and kanamycin. The next day, colonies were restreaked and incubated at 37°C. Primers used for pKD4 amplification and colony screening are provided in Supplemental Table 1.

### RT-qPCR

Overnight cultures were diluted 1:100 in 3 mL LB containing 1nM aTc. Cells were further grown for 2h before RNA extraction using Direct-zol™ RNA MiniPrep (Zymo Research) followed by DNAse treatment using TURBO DNA-free Kit (Thermo Fisher Scientific). All RNAs were reverse transcribed into cDNA using the Transcriptor First Strand cDNA Synthesis Kit using 500 ng RNA (Roche). qPCR was performed with the FastStart Essential DNA Probes master mix (Roche) in a LightCycler 96 (Roche) following the manufacturer’s instructions. qPCR was performed in two technical replicates and three biological replicates. Relative gene expression was computed on LightCycler 96 software (Roche) using the ΔΔCq method (Schmittgen and Livak 2008) after normalization by 5S rRNA (*rrsA*). Primers are listed in Supplemental Table 9.

## Data Access

The screen results are provided as Supplemental Tables 2-4. Other relevant data supporting the findings of the study are available in this article and its Supplemental Information files, or from the corresponding author upon request.

## Acknowledgements

We thank Alicia Calvo-Villamañán and Olaya Rendueles-Garcia for helpful advice as well as Antoine Decrulle for providing phage 186cIts. This work was supported by the European Research Council (ERC) under the Europe Union’s Horizon 2020 research and innovation program (grant agreement No [677823]), by the French Government’s Investissement d’Avenir program and by Laboratoire d’Excellence ‘Integrative Biology of Emerging Infectious Diseases’ (ANR-10-LABX-62-IBEID); F.R is supported by a doctoral scholarship from Ecole Normale Supérieure.

